## Supplemental Figures and Tables for "Small RNAs from Mitochondrial Genome Recombination Sites are Incorporated into *T. gondii* Mitoribosomes"

### Tetzlaff et al. - Supplemental Figures and tables

### Supplemental figures

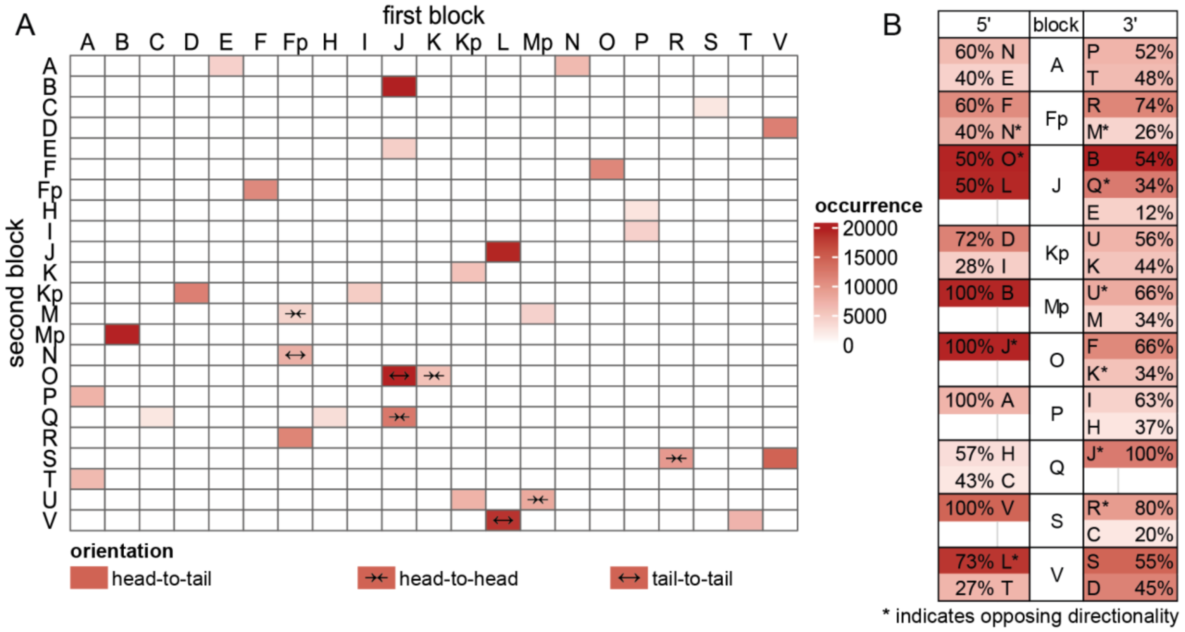

**Figure S1: Counting of *T. gondii* mitochondrial sequence block organization.**

Mitochondrial sequence blocks [1] were annotated on *T. gondii* mitochondrial ONT reads with at least 75% similarity. The annotations and their respective directionality were extracted and all two-block combinations, separated by less than 10 bases, were quantified using a custom R script. Pairs occurring less than 50 times were identified as false-positives and consequently removed from the figure.

(A) Unordered heatmap of two-block combinations. 'Head-to-tail' denotes that the 3' end of the first block is followed by the 5' end of the second block, while 'head-to-head' indicates that the 3' end of the first block is followed by the 3' end of the second block. 'Tail-to-tail' indicates a proximity of 5' ends.

(B) Block frequencies for all blocks neighbored by at least three other blocks. The colors represent the absolute number of respective neighbors, while the relative number of identified 5' and 3' neighbors is presented in percentage values. Asterisks are used to highlight opposing annotated directionalities.

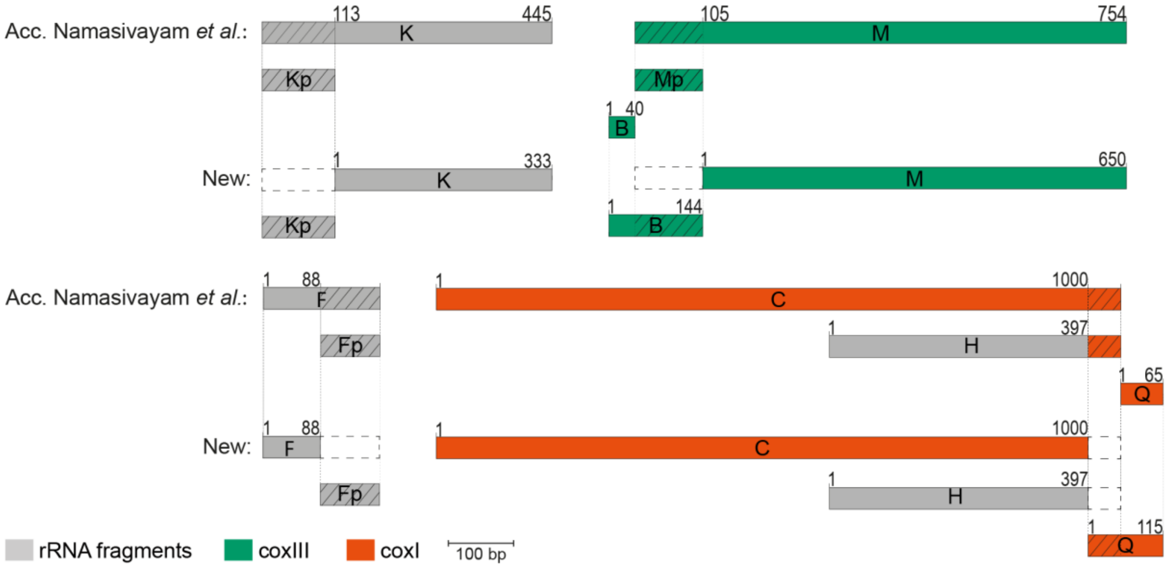

**Figure S2: Proposed changes to the sequence blocks previously annotated by Namasivayam et al. 2021 [1].**

Upper panel and lower panel left side: the blocks Kp, Mp, and Fp (p for partial) were previously defined as truncated versions of K, M and F [1]. However, the blocks Fp, Kp, and Mp frequently occur separately in the mitochondrial genome. We therefore treated Fp, Kp and Mp as separate blocks and have shortened the blocks F, K and M accordingly. This leads to a more consistent block definition without overlapping blocks and was important for our quantification of block combinations (Fig. S1). The sequence of block Mp was merged with block B since block B did not appear as an independent block in our block combination analysis (Fig S1, Tab S4). Lower panel right: The blocks C and H terminate with a short duplication of the *coxI* coding sequence. We propose to add this sequence to block Q, which is in all cases following C and H, respectively.

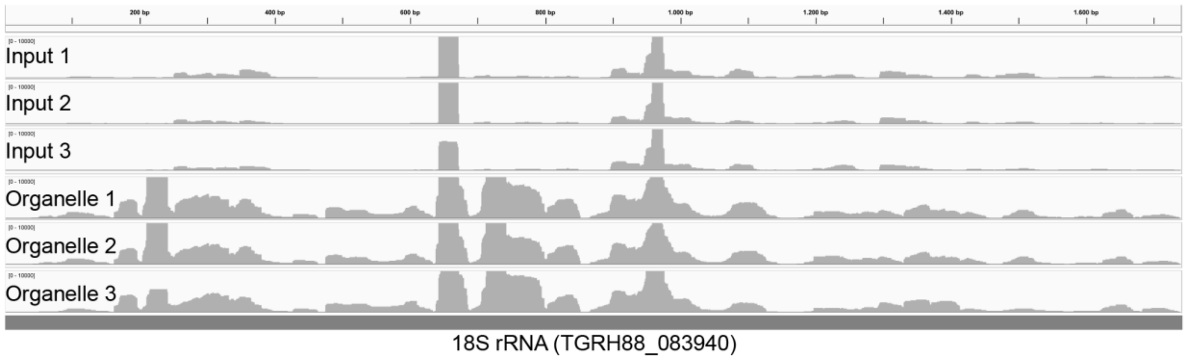

**Figure S3: sRNA sequence coverage of an 18S rRNA gene.**

Mapping of reads from input and enriched sRNA libraries to an 18S rRNA gene demonstrates that the enriched libraries have a much higher coverage than input libraries. This suggests that the enrichment procedure generates cytosolic rRNA fragments that are co-purified, for example from partially digested, ER-membrane bound ribosomes. This pattern was observed for all cytosolic rRNAs (not shown). Since the benzonase does not degrade the cytosolic components entirely but creates a large number of smaller fragments, more reads for cytosolic rRNA were eventually found in the enriched RNA library than in the input library.

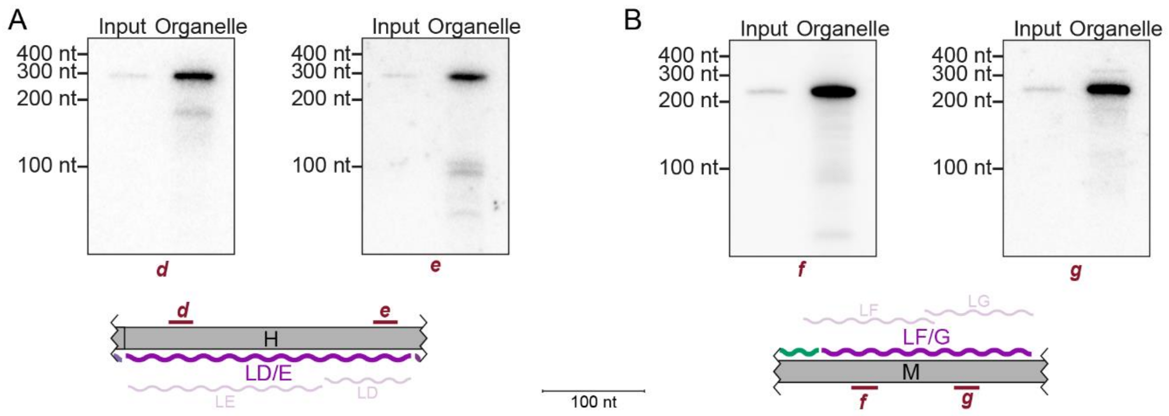

**Figure S4: Detection of LSUD/E and LSUF/G transcripts.**

(A) RNA gel blot hybridization of LSUE (left) and LSUD (right), which are located side-by-side on block H. LSUE and LSUD were both detected at a size of 300 nt with a much stronger signal in the organellar enriched RNA fraction than in the total input RNA. This length corresponds to a co-transcript of LSUE and LSUD. Signals for smaller RNAs were detected (probe d= ~200 nt, probe e = 100 nt), but to a much lesser extent than the longest fragment. Bottom: Schematic representation of the position of the LSUE (LE) and LSUD (LD) transcripts (light coloured wiggle lines) and the single LD/E transcript (dark colored wiggle line) relative to genomic block H. Green wiggle-line: end of *coxIII* transcript. Red bars with lower case letters indicate positions of probes used for detecting RNAs in RNA gel blot hybridizations.

(B) Same as in (A), but for the LSUF and LSUG RNAs. LSUF and LSUG were detected at a size of about 240 nt again with a much stronger signal in the organellar enriched RNA fraction and negligible amounts of smaller RNAs.

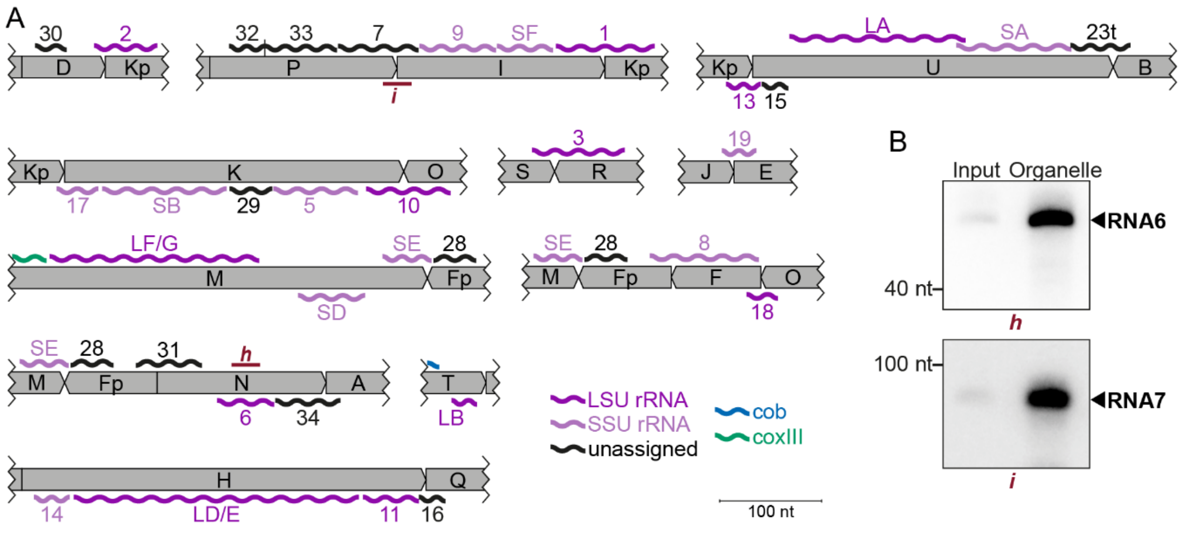

**Figure S5: Overview of non-coding RNAs identified in *T. gondii* mitochondria.**

(A) Schematic representation of the position of all non-coding transcripts in the *T. gondii* genome. Transcripts are shown as wiggle lines and are named according to prior nomenclature [2]. Assignment of transcripts to the large and the small rRNA are indicated by color. Red bars with lower case letters indicate positions of probes used for detecting RNAs in RNA gel blot hybridizations in (B). L/S = LSU/SSU = large/small subunit of the ribosome.

(B) RNA gel blot hybridization of RNA6 and RNA7 using RNA extracts from input and organelle enriched fractions of the organelle enrichment protocol shown in Fig. 1.

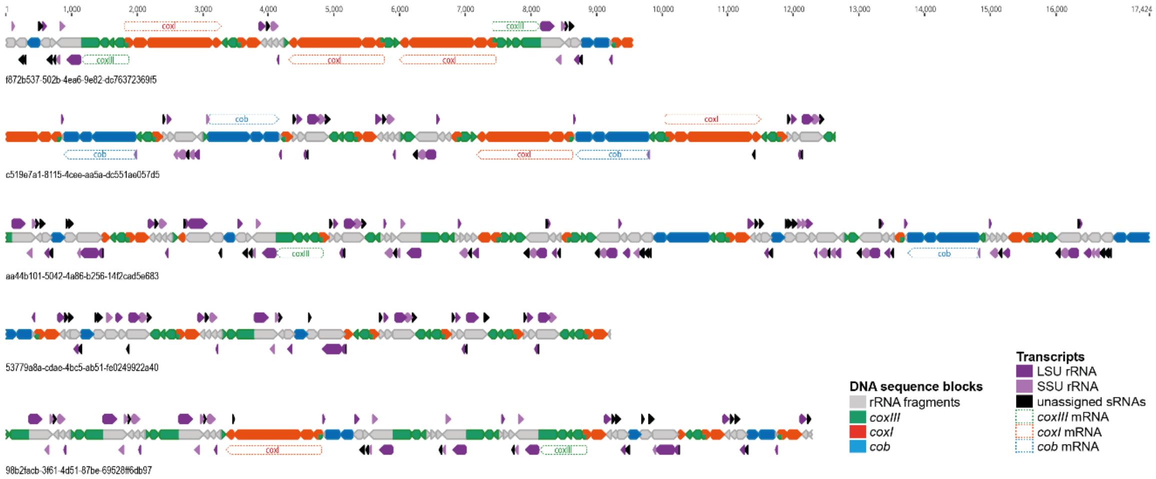

**Figure S6: Mapping of small RNAs onto exemplary ONT sequencing reads.**

ONT sequencing reads are drawn to scale with block annotations indicated by a color code. Numbers below reads are unique identifiers in the published sequence read archive (acc. no PRJNA978626 at SRA). The blocks are shown as arrows to indicate the direction of blocks as previously defined [1]. Sequence blocks located on the ends of reads are incomplete. sRNAs are shown above and below the ONT depending to which strand they belong. According to their homology to rRNAs they are assigned either to the small (SSU) or the large (LSU) subunit, which is indicated by arrow color. Potential full-length mRNAs for the three reading frames are shown as well. Please note that the non-coding sequence blocks are almost fully covered by small RNA species and that full-length mRNAs can be produced from multiple locations.

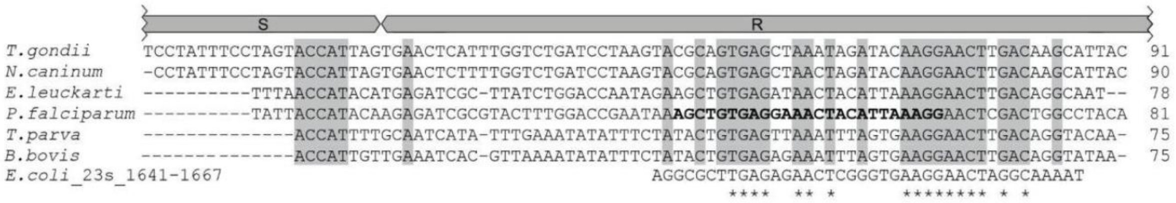

**Figure S7: Alignment of RNA3 sequences from different apicomplexan species.**

Alignment of apicomplexan RNA3 sequences using Clustal Omega (https://www.ebi.ac.uk/Tools/msa/clustalo/). Sequence excerpts from *E. coli* corresponding to sections of the alignment were added manually based on previous analyses [2]. Nucleotide positions conserved in all apicomplexans are shaded in gray. The residues in the *P. falciparum sequence* suggested to replace sequence elements in the *E. coli* 23S rRNA structure [2] are marked in bold. Residues conserved between apicomplexans and *E. coli* are marked with asterisks.

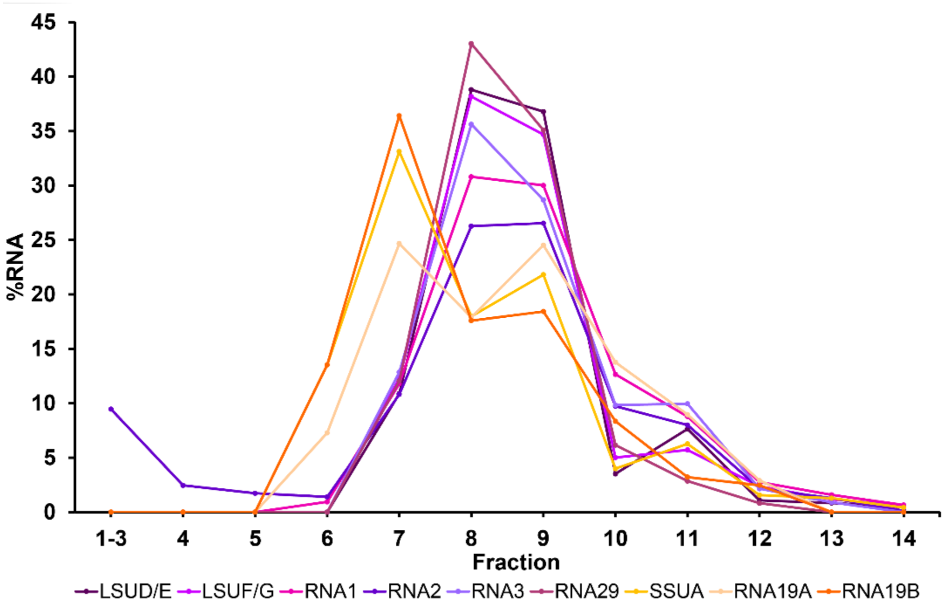

**Figure S8: Migration patterns of mitochondrial sRNAs in sucrose density gradients under $Mg^{2+}$-depleted conditions**

The distribution of different small RNAs in sucrose density gradients, conducted in absence of $Mg^{2+}$ and presence of 10 mM EDTA, was analysed by RNA gel blot hybridizations (Fig 6). Signals were quantified and visualized for each small RNA as percentage of RNA per fraction. Small RNAs assigned to the ribosomal SSU are visualized in orange hues, while those allocated to the ribosomal LSU are shown in lilac shades. Signals for SSU rRNAs start in fraction 6 and peak in fraction 7 whereas LSU signals start in fraction 7 and peak in fraction 8 and 9.

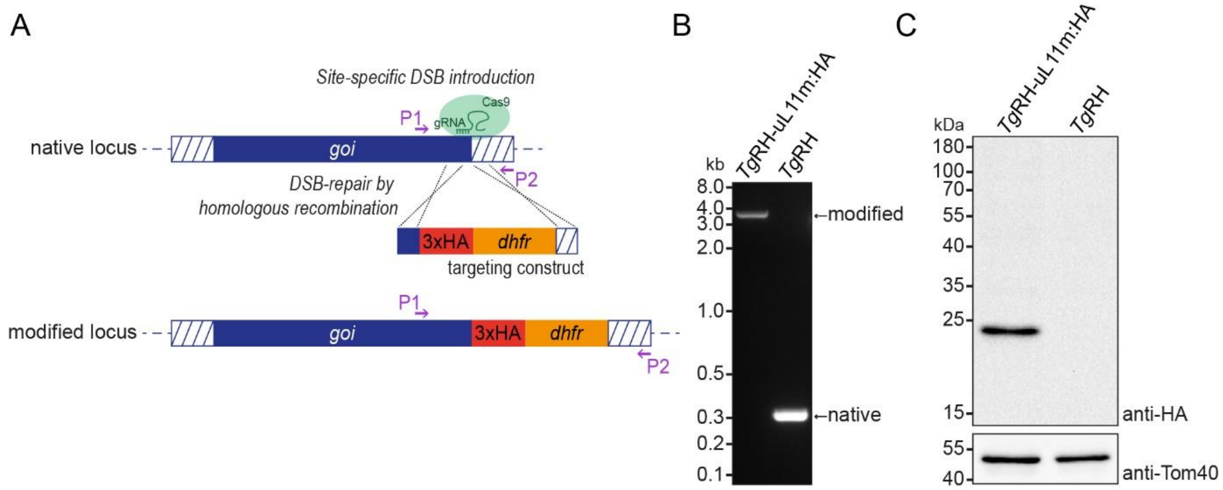

**Figure S9: Generation of a tagged ribosomal protein TgRH-uL11mHA.**

(A) The 3' replacement sequence consists of homologous regions to the gene of interest, here *Tg*RH-uL11mHA, and includes the HA-tag in frame with the reading frame at the native locus as well as the *dhfr* marker gene. This is introduced via homologous recombination induced by a CRISPR/Cas-mediated double-stranded break in the target sequence to yield the modified locus. (B) A PCR using primers P1 and P2 (see A) on the parental strain *Tg*RH and the modified strain *Tg*RH-uL11mHA demonstrates successful modification of the target locus. (C) An immunoblot analysis of *Tg*RH-uL11m:HA-expressing parasites and parental strain *Tg*RH parasites was performed using an anti-HA antibody and as a loading control, an anti-Tom40 antibody.

### Supplemental tables

**Table S1: ONT sequencing mapping statistics on nuclear and organellar genomes for data obtained in this study and data previously published [1].**

| | total length (nt) | | | |
|---|---|---|---|---|
| | **this study RH** | **this study mt reads >1000nt** | **Namasivayam et al. ME49** | **Namasivayam et al. RH Δuprt** |
| *raw reads total* | *1.060.579.347* | | | |
| *others* | *184.244.447* | | | |
| *apicoplast* | *22.322.153* | | | |
| mt | 78.540.610 | 48.661.147 | 699.177 | 1.985.929 |
| nucleus | 775.472.137 | 805.351.600 | 313.953.499 | 4.309.669.177 |
| mt + nucleus | 854.012.747 | 854.012.747 | 314.652.676 | 4.311.655.106 |
| **% mt** | **9,20** | **5,70** | **0,22** | **0,05** |
| | **#reads** | | | |
| | **this study RH** | **this study mt reads >1000nt** | **Namasivayam et al. ME49** | **Namasivayam et al. RH Δuprt** |
| *raw reads total* | *2.080.000* | | | |
| *others* | *299.025* | | | |
| *apicoplast* | *8.152* | | | |
| mt | 86.761 | 23.693 | 269 | 779 |
| nucleus | 1.686.062 | 1.749.130 | 43.392 | 765.200 |
| mt + nucleus | 1.772.823 | 1.772.823 | 43.661 | 765.979 |
| **% mt** | **4,89** | **1,34** | **0,62** | **0,10** |

**Table S2: Read length distribution of *T. gondii* mitochondrial ONT reads.**

| Read length (nt) | Number of reads |
|---|---:|
| 0 - 150 | 103 |
| 150 - 500 | 38,343 |
| 500 - 1,000 | 24,643 |
| 1,000 - 2,000 | 15,402 |
| 2,000 - 3,000 | 4,710 |
| 3,000 - 4,000 | 1,864 |
| 4,000 - 5,000 | 860 |
| 5,000 - 6,000 | 399 |
| 6,000 - 7,000 | 183 |
| 7,000 - 8,000 | 118 |
| 8,000 - 9,000 | 53 |
| 9,000 - 10,000 | 38 |
| >10,000 | 45 |
| **total** | 86,761 |

**Table S3: Mitochondrial sequence block frequencies in ONT DNA sequencing data.**

| Block | Number |
|---|---|
| A | 19794 |
| B | 21737 |
| C | 3138 |
| D | 14076 |
| E | 5703 |
| F | 12089 |
| Fp | 21286 |
| H | 3572 |
| I | 5637 |
| J | 45849 |
| K | 6629 |
| Kp | 19398 |
| L | 22564 |
| M | 4947 |
| Mp | 21991 |
| N | 8393 |
| O | 21814 |
| P | 9940 |
| Q | 14275 |
| R | 12662 |
| S | 18294 |
| T | 8780 |
| U | 9431 |
| V | 33337 |

**Table S4: Sequence block combinations identified in *T. gondii* mitochondrial ONT reads using a custom R script.**

Combinations that were found less than 50-times are considered false-positives and shown in grey.

| 1st block | direction | 2nd block | direction | n | orientation |
|---|---|---|---|---|---|
| J | forward | B | forward | 19622 | head-tail |
| J | reverse | O | forward | 19528 | tail-tail |
| B | forward | Mp | forward | 19378 | head-tail |
| L | forward | J | forward | 19307 | head-tail |
| L | reverse | V | forward | 18377 | tail-tail |
| V | forward | S | forward | 14445 | head-tail |
| J | forward | Q | reverse | 12391 | head-head |
| V | forward | D | forward | 11757 | head-tail |
| D | forward | Kp | forward | 11741 | head-tail |
| Fp | forward | R | forward | 11400 | head-tail |
| O | forward | F | forward | 11044 | head-tail |
| F | forward | Fp | forward | 10765 | head-tail |
| R | forward | S | reverse | 9320 | head-head |
| Mp | forward | U | reverse | 7858 | head-head |
| Fp | reverse | N | forward | 7301 | tail-tail |
| Kp | forward | U | forward | 6969 | head-tail |
| A | forward | P | forward | 6957 | head-tail |
| T | forward | V | forward | 6858 | head-tail |
| N | forward | A | forward | 6345 | head-tail |
| A | forward | T | forward | 6320 | head-tail |
| K | forward | O | reverse | 5680 | head-head |
| Kp | forward | K | forward | 5447 | head-tail |
| I | forward | Kp | forward | 4599 | head-tail |
| J | forward | E | forward | 4345 | head-tail |
| P | forward | I | forward | 4274 | head-tail |
| E | forward | A | forward | 4248 | head-tail |
| Mp | forward | M | forward | 4078 | head-tail |
| Fp | forward | M | reverse | 3989 | head-head |
| H | forward | Q | forward | 3060 | head-tail |
| P | forward | H | forward | 2525 | head-tail |
| C | forward | Q | forward | 2293 | head-tail |
| S | forward | C | forward | 2287 | head-tail |
| J | reverse | R | forward | 45 | tail-tail |
| V | forward | V | reverse | 18 | head-head |
| Fp | forward | O | forward | 14 | head-tail |
| D | forward | A | forward | 10 | head-tail |
| D | forward | D | reverse | 6 | head-head |
| I | forward | S | forward | 6 | head-tail |
| S | forward | E | forward | 5 | head-tail |
| Kp | reverse | Kp | forward | 5 | tail-tail |
| J | reverse | J | forward | 5 | tail-tail |
| D | forward | I | reverse | 3 | head-head |

| | | | | | |
|---|---|---|---|---|---|
| S | forward | M | forward | 3 | head-tail |
| S | reverse | S | forward | 3 | tail-tail |
| B | forward | Mp | reverse | 2 | head-head |
| Q | forward | A | forward | 2 | head-tail |
| T | forward | L | forward | 2 | head-tail |
| V | forward | Kp | forward | 2 | head-tail |
| Q | forward | B | forward | 2 | head-tail |
| S | reverse | C | forward | 2 | tail-tail |
| S | reverse | D | forward | 2 | tail-tail |
| P | reverse | P | forward | 2 | tail-tail |
| A | forward | E | reverse | 1 | head-head |
| B | forward | V | forward | 1 | head-tail |
| F | forward | O | reverse | 1 | head-head |
| I | forward | J | forward | 1 | head-tail |
| I | forward | P | reverse | 1 | head-head |
| J | forward | Mp | reverse | 1 | head-head |
| Kp | forward | Kp | reverse | 1 | head-head |
| L | forward | O | forward | 1 | head-tail |
| Mp | forward | C | forward | 1 | head-tail |
| P | forward | P | reverse | 1 | head-head |
| Q | forward | A | reverse | 1 | head-head |
| S | forward | S | reverse | 1 | head-head |
| S | forward | F | reverse | 1 | head-head |
| U | forward | J | reverse | 1 | head-head |
| Q | reverse | Q | forward | 1 | tail-tail |
| Kp | reverse | K | forward | 1 | tail-tail |
| B | reverse | Mp | forward | 1 | tail-tail |
| A | reverse | A | forward | 1 | tail-tail |
| L | forward | V | forward | 1 | head-tail |
| P | reverse | T | forward | 1 | tail-tail |
| L | reverse | L | forward | 1 | tail-tail |
| L | reverse | Mp | forward | 1 | tail-tail |
| J | forward | Mp | forward | 1 | head-tail |
| P | forward | K | forward | 1 | head-tail |
| J | reverse | S | forward | 1 | tail-tail |
| Fp | reverse | Fp | forward | 1 | tail-tail |
| I | forward | Fp | forward | 1 | head-tail |
| O | reverse | Q | forward | 1 | tail-tail |
| E | reverse | V | forward | 1 | tail-tail |
| Mp | forward | E | forward | 1 | head-tail |
| J | forward | F | forward | 1 | head-tail |
| C | reverse | U | forward | 1 | tail-tail |

**<u>Table S5: Fraction of reads containing full-length open reading frames.</u>**

| | ORF length [nt] | #reads longer than ORF length | #reads containing full-length ORF | % reads containing ORF |
|---|---|---|---|---|
| **coxI** | 1,476 | 12,099 | 1,337 | **11.1** |
| **coxIII** | 745 | 28,503 | 1,463 | **5.1** |
| **cob** | 1,107 | 18,149 | 1,560 | **8.6** |

**Table S6: Mapping statistics of the RNA-seq data on the different genomes/sequence blocks.** After filtering the raw reads against the nuclear rRNA genes, the remaining reads were mapped against the three subgenomes of *T. gondii* RH-88.

| Library | reads after rRNA filtering | Nuclear genome[1] | Apicoplast genome[2] | Mitochondrial sequence blocks[3] | Mitochondrial sequence blocks and combinations[4] |
|---|---|---|---|---|---|
| Input1 | 14.402.531 | 8.380.729 | 27.627 | 4.182.533 | 4.814.603 |
| Input2 | 9.207.557 | 5.367.352 | 21.958 | 2.481.756 | 2.898.989 |
| Input3 | 16.001.184 | 7.791.619 | 34.156 | 5.866.849 | 6.905.255 |

1 from strain RH-88, accession number GCA_019455545.1

2 accession number CM033583.1

3 individual blocks according to (Namasivayam et al. 2021; MN077088.1- MN077111.1)

4 All block combinations identified here as described in Fig. 2

**Table S7: List of mitochondrial non-coding RNAs identified by sRNA sequencing.**

| Accession | Block | Assigned Name[1] | Position from RNA sequencing [Position previous annotation] | Length | assigned rRNA region[2] | Only found in cyst-forming Eucoccidians |
|---|---|---|---|---|---|---|
| MN077107.1 | U | **RNA15** | complement (10-36) | 27 | | |
| | | LSUA | 37 – 209 [37 – 209] | 173 | LSU1 | |
| | | SSUA | 201 – 313 [204 – 314] | 113 | SSU4 | |
| OR086911 | K | SSUB | complement (38-159) [complement (38-156)] | 122 | SSU6 | |
| | | **RNA29** | complement (163-204) | 42 | | + |
| | | **RNA5** | complement (206-288) | 83 | SSU9 | |
| MN077095.1 | I | **RNA9** | 23 – 97 | 75 | SSU8 | |
| | | SSUF | 99 – 153 [complement (98-156)] | 55 | SSU12 | |
| OR086913 | H | **RNA14** | complement (13-47) | 35 | SSU1 | |
| | | LSUE | complement (52-331) [complement (54-244; 247-331)] | 280 | LSU9 | |
| | | LSUD | | | LSU8 | |
| | | **RNA11** | complement (336-390) | 55 | LSU5 | |
| OR086912 | M | LSUF | 280 – 485 [262-390; 381 – 487] | 206 | LSU11 | |
| | | LSUG | | | LSU12 | |
| | | SSUD | complement (523-588) [complement (524-588)] | 66 | SSU10 | |
| MN077106.1 | T | LSUB | complement (246-271) [complement (247-271)] | 25 | LSU3 | |
| MN077109.1 | Fp | **RNA28** | complement (44-85) | 42 | | + |
| MN077102.1 | P | **RNA32** | 20 - 54 | 35 | | + |
| | | **RNA33** | 55 - 125 | 71 | | + |
| MN077100.1 | N | **RNA6** | complement (60-115) | 56 | LSU15 | |
| MN077091.1 | D | **RNA30** | 14 – 44 | 31 | | + |
| MN077102.1 + MN077095.1 | P + I | **RNA7** | Block P 127-184 Block I 1-21 | 79 | | |
| MN077101.1 + OR086910 | O + F | **RNA18** | Block O 69-86 Block F 1-13 | 31 | LSU14 | |
| MN077101.1 + MN077097.1 | O + K | RNA10 | Block O 47-86 Block K complement (291-333) | 83 | LSU13 | |
| OR086912 + MN077109.1 | M + Fp | SSUE | Block M 606-650 Block Fp complement (89-91) | 48 | SSU11 | |
| MN077109.1 + MN077100.1 | Fp + N | **RNA31** | Block Fp complement (1-20) Block N 1-45 | 65 | | + |
| MN077109.1 + OR086910 | Fp + F | RNA8 | Block Fp complement (1-20) Block F complement (2-88) | 107 | SSU5 | |
| MN077091.1 + MN077110.1 | D + Kp | **RNA2** | Block D 72-82 Block Kp 1-52 | 63 | LSU2 | |
| MN077095.1 + MN077110.1 | I + Kp | **RNA1** | Block I 157-204 Block Kp 1-48 | 96 | LSU6 | |

| | | | | | |
|---|---|---|---|---|---|
| MN077107.1<br>+<br>MN077110.1 | U<br>+<br>Kp | **RNA13** | Block U complement (1-8)<br>Block Kp complement (87-112) | 34 | LSU10 |
| OR086911<br>+<br>MN077110.1 | K<br>+<br>Kp | **RNA17** | Block K complement (1-33)<br>Block Kp complement (105-112) | 41 | SSU3 |
| OR086915<br>+<br>OR086913 | Q<br>+<br>H | **RNA16** | Block Q complement (1-18)<br>Block H complement (390-397) | 26 | |
| MN077107.1<br>+<br>OR086916 | U<br>+<br>B | **RNA23t** | Block U 312-354<br>Block B complement (127-144) | 61 | |
| MN077088.1<br>+<br>MN077100.1 | A<br>+<br>N | **RNA34** | Block A complement (1-13)<br>Block N complement (117-166) | 63 | |
| MN077105.1<br>+<br>MN077104.1 | S<br>+<br>R | **RNA3** | Block S 184-205<br>Block R complement (17-85) | 91 | LSU7 |
| MN077096.1<br>+<br>MN077092.1 | J<br>+<br>E | **RNA19** | Block J = 74-85<br>Block E = 1-22 | 34 | SSU7 |

1 RNAs in bold have not been previously predicted based on sequence similarities [1]
2 Numbers specify the linear order of fragments relative to conventional rRNA as suggested by Feagin *et al.* 2012 [2] (LSU: ribosomal large subunit, SSU: ribosomal small subunit)

**Table S8: Overview of mitochondrial non-coding RNAs identified in *P. falciparum* and *T. gondii***

| | *P. falciparum** | | *T. gondii* | |
|---|---|---|---|---|
| | number of fragments | nt total | number of fragments | nt total |
| LSU rRNA | 15 | 1233 | 12[$] | 1193 |
| SSU rRNA | 12 | 804 | 11 | 845 |
| unassigned sRNAs shared[1] | 4 | 205 | 4 | 193 |
| unassigned sRNAs unique[2] | 8 | 350 | 7 | 349 |

* based on Feagin *et al.,* 2012 and Hillebrand *et al.,* 2018

[$] Please note that LSUE and D as well as LSUF and G were shown to be single rRNAs here, thus reducing the number of fragments in *T. gondii*

[1] sRNAs not assigned to a region of ribosomal RNA, found in *P. falciparum* and *T. gondii*

[2] sRNAs not assigned to a region of ribosomal RNA, only found in *P. falciparum* or *T. gondii*

**Table S9: List of oligonucleotides used in this study**

| Name | Sequence (5' to 3') | ID* |
|---|---|---|
| RNA17 probe | GACTTATTAAACCAGCCTGGGATCA | a |
| RNA29 probe | GTTCTAATTCCCCGTGGTAAACACAGTC | b |
| RNA5 probe | GTGTCATAGGAGATATACTCTATAATTAG | c |
| LSUE probe | GCCATCTCGATCCTCATATTCAAA | d |
| LSUD probe | GCGGTATCTAGTGTTGATGTACAA | e |
| LSUF probe | GAACTCCCAATATTATCTGACCTGTT | f |
| LSUG probe | GTATTACATCTGACGGTGAACTATC | g |
| RNA6 probe | GTCTCTCAATAACTAGCTGAGTGCTTG | h |
| RNA7 probe | GTTTCATAAATACAATCAGTGAAAGCTCTT | i |
| RNA1+RNA2 probe | GTTTGTACCTACTTGACTCCTCAGTTTAAG | j |
| RNA34 probe | GCATTTCCTATGCTCCTTAACATCACAGC | k |
| RNA19 probe | GTTCTCGAAACCATGCTAACACAATAG | l |
| RNA3 probe | GCTCACTGCGTACTTAGGATCAGACCAA | m |
| SSUA probe | GACTCTTCTATAGTTTAACCGCTACTG | |
| SSUD probe | GGCGCTTAATAACGATTCCGTC | |
| Cob RACE PCR primer | CCCCAGAAACTCATTTGTCCCC | |
| Rumsh1 RACE PCR primer | TGATCCAACCGACGCGAC | |
| Rumsh 5' RACE linker | GUGAUCCAACCGACGCGACAAGCUAAUGCAAGANNN (RNA) | |
| cox3 qPCR fwd | CAGTACCATCGTACTGGGAGTAATC | |
| cox3 qPCR rev | GATAGACCTAAGTATTCCGTACAGAC | |
| cob qPCR fwd | CGCGCTTAAAGTTGCCTTTTATC | |
| cob qPCR rev | GCTCGAATCTCAGAAAGTAAACC | |
| cox1 qPCR fwd | GAGTTATACAGTTCTGGTTCGC | |
| cox1 qPCR rev | CACTACCAAATTCAGCACAAATAC | |
| LSUF/G qPCR fwd | GTCGGTGCTATCTTGGGTTTC | |
| LSUF/G qPCR rev | CCTGTTATCCCCGGCGTACCTTAC | |
| ESR1 qPCR fwd | CCAGATGGTCAGTGCCTTGT | |
| ESR1 qPCR rev | CAAATCCACAAAGCCTGGCA | |
| 3x HA amplification fwd – (+HindIII restriction site) | TATAAAGCTTGGTGGAGGTAGCGGTGGTGGAAGTT ACCCGTACGACGTCCCG | |
| 3x HA amplification rev – (+NsiI restriction site) | TATAATGCATATTATGCGCAGGCATAATCTGGAACA TCGTAAGG | |
| Q5 mutagenesis gRNA fwd Tgurpl11m tagging | GTCACAGTCACTTCTTTGTGGGTTTTAGAGCTAGA AATAGC | |
| Q5 mutagenesis rev Tgurpl11m tagging | AACTTGACATCCCCATTTAC | |
| flank fwd Tgurpl11m tagging | AAGACATGCGTCAGAGAAAGAAAGCAGCGAAGCGA GCGGCCACAAAGAAGGGTGGAGGTAGCGGTGGTG GAAGT | |
| flank rev Tgurpl11m tagging | TCTTGAAACAGACGGGAAGGCAAAAAGAACAAAAA AACCACACTTCACAGTGCAGGGCTCTAGAACTAGT | |

| | | |
|---|---|---|
| | GGATCG | |
| genotyping fwd Tgurpl11m tagging | TTGCCTGCGTACCACAAGC | P1 |
| genotyping rev Tgurpl11m tagging | CAGGAGAAAGCCAAGCGGA | P2 |

* lowercase letter used as probe IDs or numbers used as primer IDs in figures, respectively
